# Phosphatidylserine-Based Liposomes Encapsulating DMX-5804 Protect Against Doxorubicin-Induced Cardiotoxicity

**DOI:** 10.64898/2026.02.12.705423

**Authors:** Jessica Tetterton-Kellner, Brian C. Jensen, Juliane Nguyen

**Affiliations:** Division of Pharmacoengineering and Molecular Pharmaceutics, Eshelman School of Pharmacy, University of North Carolina at Chapel Hill, Chapel Hill, NC 27599, USA; McAllister Heart Institute, University of North Carolina, Chapel Hill, NC 27599, USA; Department of Medicine, Division of Cardiology, University of North Carolina, Chapel Hill, NC 27599, USA

## Abstract

Anthracycline induced cardiotoxicity is a significant problem for oncologists and cancer patients. The leading cause of non-cancer death in cancer patients and survivors is heart failure, which is frequently attributed to the exposure to chemotherapeutics like anthracyclines. The most notorious of these chemotherapies is doxorubicin, which causes cardiac contractile dysfunction that in some cases is irreversible. In this study, we report the development of NanoDMX, a phosphatidylserine-containing liposomal formulation of DMX5804, a small molecule inhibitor of MAP4K4, and demonstrate that its administration prevents doxorubicin-induced left ventricular dysfunction in mice. Additionally, we demonstrate that DMX-5804 protects cardiomyocytes *in vitro* through a combination of mechanisms outside of the expected route of suppressing the JNK pathway. Overall, we demonstrate that the use of NanoDMX, a novel liposomal system using both DMX-5804 and phosphatidylserine, can prevent the damage induced by doxorubicin over the course of a single high dose *in vivo* model.

## INTRODUCTION

Chemotherapy-induced cardiotoxicity represents a significant and increasingly recognized consequence of modern cancer therapy.^1^ As oncologic treatments continue to improve in efficacy and extend patient survival, a growing number of patients are exposed to agents with well-documented cardiotoxic potential. While the success of current chemotherapy marks a major milestone in oncology, it simultaneously highlights the urgent need for comprehensive, long-term surveillance of survivorship outcomes. Notably, cardiovascular disease has emerged as the leading cause of non-cancer-related mortality among cancer survivors, a trend largely driven by the cumulative cardiotoxic effects of chemotherapeutics agents administered throughout treatment regimens.^2^

Currently, dexrazoxane remains the only FDA-approved drug for preventing or mitigating chemotherapy-induced cardiotoxicity. As an iron chelator, it reduces oxidative myocardial damage by reducing the formation of iron-anthracycline complexes. However, its clinical use is limited by several factors. Dexrazoxane is associated with adverse effects that resemble those of doxorubicin, including myelosuppression and gastrointestinal symptoms, as well as concerns about potential interference with chemotherapeutic efficacy. Although recent studies have reaffirmed its cardioprotective benefit, particularly in pediatric patients with acute myeloid leukemia (AML), dexrazoxane remains contraindicated in pregnancy^3^, and its mechanism of action remains incompletely understood. Interestingly, it has been identified as a topoisomerase IIB inhibitor^4^, the same target through which doxorubicin induces DNA damage in cardiomyocytes, raising questions about how dexrazoxane selectively protects the heart while preserving antitumor activity. These limitations collectively highlight the need for novel, mechanistically distinct, and safer cardioprotective therapeutics.

However, the development of such therapeutics is further complicated by the multifactorial mechanisms underlying doxorubicin-induced cardiotoxicity and the incomplete understanding of the many mechanisms that contribute to cardiomyocyte injury following doxorubicin exposure. Among these mitochondria-induced apoptosis has emerged as a central mediator.^5^ Doxorubicin preferentially accumulates within mitochondria due to their high cardiolipin content, where it disrupts electron transport chain function, enhances reactive oxygen species (ROS) generation, and triggers cytochrome c release, ultimately activating intrinsic apoptotic cascades.^6^ In addition to mitochondria-dependent apoptosis, other regulated cell death pathways including necroptosis^7^ and ferroptosis^8^ contribute to doxorubicin-induced myocardial injury. Furthermore, doxorubicin’s canonical mechanism involving topoisomerase IIβ–mediated DNA amplifies mitochondrial dysfunction and oxidative stress, compounding cardiomyocyte vulnerability.^9^

Given the high mitochondrial density of cardiomyocytes compared to other cell types, the mitochondrial protection in cardiomyocytes represents a promising therapeutic strategy against anthracycline-induced cardiotoxicity.^10^ In parallel, modulation of stress-activated signaling pathways has emerged as an alternative means to preserve cardiomyocyte viability. Among these, MAP4K4 has been implicated in the regulation of oxidative stress, apoptosis, and inflammation within cardiac tissue, making it an attractive target for cardioprotection.^11^

DMX-5804, developed to inhibit MAP4K4, was previously investigated for its potential to treat myocardial infarctions.^12^ However, its clinical translation has been hampered by relatively poor solubility and limited efficacy in large-animal models of myocardial infarction. To improve the solubility of the drug, a prodrug was created by researchers. However, upon the successful generation of a soluble prodrug of DMX-5804, known as DMX-10001, high plasma concentrations of DMX-5804 were observed, but no reduction in infarct size was observed in a large animal model of myocardial infarction.^13^ Building upon these observations, we sought to explore whether nanocarrier-based delivery could enhance the therapeutic potential of MAP4K4 inhibition in the context of doxorubicin-induced cardiotoxicity by improving its solubility and hence delivery efficacy.

In the present study, we developed a phosphatidylserine-enriched liposomal formulation NanoDMX, designed to deliver DMX-5804 while providing intrinsic anti-inflammatory benefits. This dual-mechanism approach aims to protect cardiomyocytes by preserving mitochondrial function and attenuating stress signaling. Through this strategy, we aim to overcome previous formulation challenges, enhance targeted delivery, and establish a foundation for next-generation cardioprotective nanotherapeutics in the context of doxorubicin-induced cardiac injury.

## MATERIALS AND METHODS

### Cell culture

H9c2 cells, a rat ventricular myoblast cell line, were purchased from ATCC (ATTC, catalog number CRL-1446). Cells were maintained in DMEM supplemented with 10% fetal bovine serum, 1% glutamine, 1% sodium pyruvate, and 1% penicillin/streptomycin in T-175 flasks. Phenol red-free media was used for assays involving fluorescence or absorbance. Media was changed every 3 days, and cells were kept below 75% confluency. Cells were started at passaged 2 upon receipt and discarded after passage 15.

### MTT Assay

Cells were seeded overnight in 96-well plates at a density of 5,000 cells per well. Black walled, clear bottom, tissue-culture-treated plates were used, and cells were allowed to adhere overnight. Cells were then treated with MAP4K4 inhibitors and doxorubicin for 24 hours prior to removing the drug-containing media and washed with PBS. MTT supplemented media was added to cells at a concentration of 2.5 mg/mL. After 4 hours of incubation in MTT treated media, media was removed, and the resulting crystals were solubilized using a mixture of DMSO and SDS. After incubating for 15 additional minutes, absorbance was read at 570 nm and 630 nm on a Molecular Devices Spectramax M2.

### TMRM Assay

Cells were seeded in black-walled, clear-bottom 96-well plates and allowed to adhere overnight. The following day, MAP4K4 inhibitors and doxorubicin were added to fresh media and incubated for 24 hours. Following a 24-hour incubation, cells were washed thoroughly to remove any free drug present in media. TMRM was added to media to a final working concentration of 50 nM and added to wells for 30 minutes at 37 °C. Following incubation, the excitation and emission was read at 548 nm and 573 nm.

### Mitotracker Red Staining

Cells were seeded overnight in 6-well plates using poly-L-lysine-coated coverslips at a density of 50,000 cells per chamber. After allowing cells to adhere overnight, MAP4K4 inhibitors and doxorubicin were added and incubated for 24 hours. Following treatment, drug-containing media was removed, and cells were washed with PBS. MitoTracker Red (500 nM) was added and cells were incubated for 30 minutes at 37°C. Cells were then fixed and permeabilized for 20 minutes at -20°C using ice cold methanol. DAPI was added at a concentration of 1 µg/mL for 10 minutes at room temperature to stain the nuclei. Cells were then imaged using Cy5 and DAPI channels and images were overlaid. 3-5 representative images were taken per chamber at different positions within the chamber. Mitochondrial density was measured via red fluorescence using ImageJ.

### Western blot analysis

Cells seeded in 6-well plates at a density of 300,000 cells per well and allowed to adhere overnight. Cells were then treated with doxorubicin and DMX-5804 for 4 or 24 hours. Unless otherwise specified, drugs were added to cells at 10 μM. After drug treatment, media was removed and cells were washed with PBS. Ice cold RIPA buffer supplemented with protease and phosphatase inhibitors was used to lyse cells. Thermo Scientific™ RIPA Lysis and Extraction Buffer (catalog no. PI89900) and Thermo Scientific™ Pierce™ Protease and Phosphatase Inhibitor Mini Tablets, EDTA-free (catalog no. PIA32961) were used to lyse cells and inhibit proteases and phosphatases. Protein concentrations were quantified using a Pierce 660 assay, and 12 µg of protein was run per lane of gel after being reduced with DTT. Invitrogen™ NuPAGE™ Bis-Tris Mini Protein Gels with a 4-12% gradient were used. Following electrophoresis and transfer onto 0.45μm PVDF membranes, membranes were blocked with 5% BSA at room temperature and subsequently incubated with primary antibodies in 2.5% BSA overnight at 4°C. The following primary antibodies were used: MAP4K4 (rabbit IgG, Proteintech catalog no. 55247-1-AP, 1:1000 dilution), phospho-MAP4K4 (Ser629) (rabbit IgG, Bioss, catalog no. BS-5491R, 1:2000 dilution), cJUN (rabbit IgG, catalog no. 1:1000 dilution), phospho-cJUN (Ser73) (rabbit IgG, Cell Signaling Technology catalog no. 3270, 1:1000), and beta-actin, HRP (rabbit IgG, Invitrogen, catalog no. PA1-183-HRP, 1:1000) dilution. The secondary antibody used was a Goat anti-Rabbit IgG HRP (Invitrogen, catalog no. 32460, 1:10,000 dilution). After primary antibody incubation overnight, blots were washed with PBST for 5 minutes three times at room temperature while rocking. Secondary antibody was added and allowed to incubate for 1 hour at room temperature while rocking, and prior to addition of ECL substrate (Thermo Scientific™ Pierce™ ECL Western Blotting Substrate, catalog no. FER32106X4), washed with PBST for 10 minutes three times at room temperature while rocking. Then ECL substrate was added and allowed to sit for 1-2 minutes prior to imaging on a Bio-Rad ChemiDoc imaging system. Biological replicates were run on three separate gels that were then cut below the 70 kDa band of ladder to probe for different targets of different sizes on the same blot.

### Doxorubicin uptake study

Cells were seeded at 5,000 cells per well in black-walled, clear-bottom 96 well plates and allowed to adhere overnight. The next day, doxorubicin and MAP4K4 inhibitors were added to the media and left to incubate for 24 hours. Afterwards, drug-containing media was removed, and cells were washed thoroughly with PBS to remove any doxorubicin that was not taken up by cells. Cells were then lysed using ethanol and 0.3M HCl, and fluorescence was read at the excitation/emission wavelengths of doxorubicin (485 and 590 nm, respectively). All drugs were tested at a 10 µM concentration. For all in vitro drug studies, solid compounds were solubilized in DMSO and diluted to ensure that the final drug-containing media did not exceed 0.5% DMSO

### qPCR

Primers were selected based on relevant literature. H9c2 cells were plated in 6-well plates at a density of 300,000 cells per well and allowed to adhere overnight. Cells were then treated with doxorubicin, DMX-5804, or a combination of both for either 4 hours or 24 hours. Following treatment, cells were washed and pelleted. RNA was extracted using the Qiagen RNeasy RNA extraction kit, with on-column DNase treatment to remove genomic DNA. RNA was quantified using a Nanodrop, and the absence of genomic DNA contamination was verified by agarose gel electrophoresis. Complementary DNA (cDNA) was synthesized using the iScript cDNA synthesis kit, and quantitative real-time PCR (qPCR) was performed using SYBR Green on a QuantStudio 6 Real-Time PCR System (Applied Biosystems). Equal amounts of cDNA were used for each qPCR reaction. Amplification specificity was confirmed by melt curve analysis following amplification, with all assays showing a single, specific melting peak and no evidence of primer-dimers or nonspecific products.

RT-qPCR data were normalized to the reference gene GAPDH. GAPDH is widely used as a control in RT-qPCR performed on H9c2 cells exposed to doxorubicin, enabling comparison with prior literature.^14,15^ Relative gene expression levels were quantified using the 2^-ΔΔCq^ method and were normalized to GAPDH. Results shown are the result of 3 biological replicates and 2 technical replicates per biological replicate. The primers used are as follows:

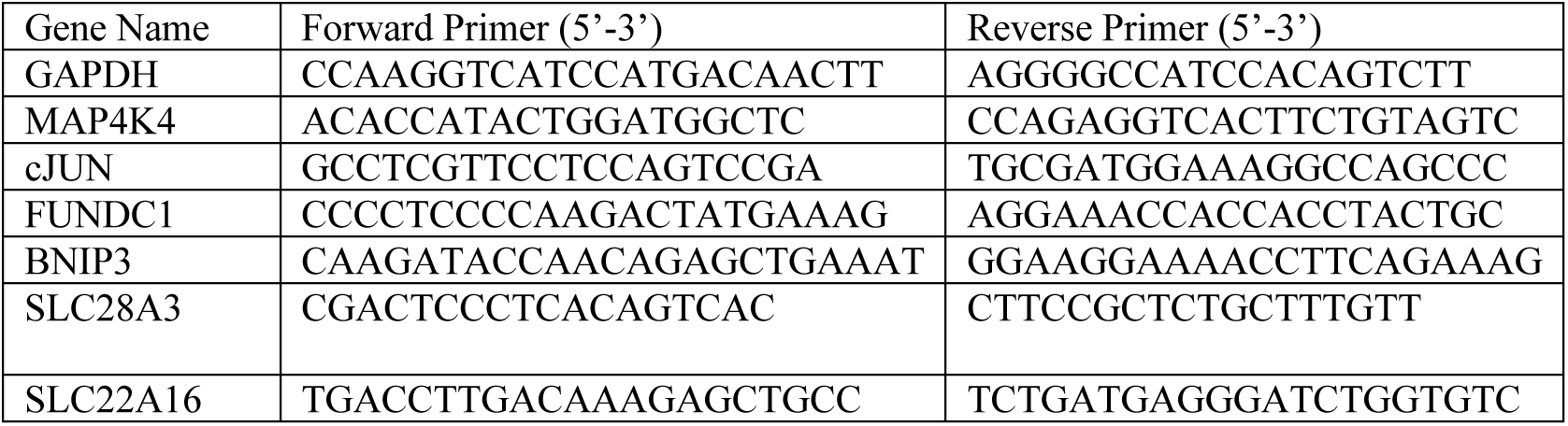

### Liposomal formulation

Liposomes were prepared using thin film hydration followed by extrusion. Lipids were purchased from Avanti Polar Lipids and solubilized in chloroform. Depending on the formulation, lipids were combined, and chloroform was evaporated using a rotary evaporator to create a thin lipid film. As DMX-5804 is a hydrophobic drug, it was dissolved in methanol and chloroform and added to the lipid solution and dried down within the thin film. The film was rehydrated using phosphate buffered saline (PBS), sonicated for 20 minutes, and extruded through polycarbonate membranes of decreasing pore sizes (1000 nm, 400 nm, 200 nm, 100 nm). Final resultant liposomes were then characterized prior to experimentation or injection via Dynamic Light Scattering (DLS). The lipids used in this study included DSPC (1,2-distearoyl-sn-glycero-3-phosphocholine), DSPS (1,2-Distearoyl-sn-glycero-3-phospho-L-serine), DSPE-PEG2000 (1,2-distearoyl-sn-glycero-3-phosphoethanolamine-N-[carboxy(polyethylene glycol)-20000), and cholesterol. DMX-5804 was solubilized in methanol prior to being dried down in the thin film of liposomes. The molar ratios of lipids tested were 60:20:20 DSPC:DSPS:Chol, and 40:40:20 DSPC:DSPS:Chol. Upon optimization, a 45% w/w ratio of DMX-5804 was loaded into the liposomes. Encapsulation efficiency was measured using UV-Vis spectroscopy. After identifying the lambda max (λ_max) of DMX-5804, liposomes encapsulating DMX-5804 were pelleted and washed to remove unencapsulated drug, and measured at λ_max before and after lysis. The encapsulation efficiency was calculated as follows: Encapsulation Efficiency (%) = Total amount of drug added/Amount of drug encapsulated in liposomes×100. Lambda max and standard curve of DMX-5804 are shown in Supplemental Figure 1.

### Animal Studies

All mice used were 10-week-old male, C57BL/6J mice from Jackson Laboratory. Animal care and experimental details were approved by the University of North Carolina Chapel Hill Institutional Animal Care and Use Committee on protocol 24-102.0. We recognize that sex (male vs female) is a biological variable that can influence physiology and response to interventions. For the purposes of this study, only male mice were used to model acute doxorubicin cardiotoxicity, a well-established approach in literature, as male mice are more susceptible than female mice to anthracycline induced cardiotoxicity.^16^ This model shows reproducible changes in the left ventricular function of mice to evaluate the cardioprotective nature of therapeutics.

### Echocardiography

To accurately measure the left ventricular ejection fraction (LVEF) of mice exposed to doxorubicin, echocardiography was performed on awake animals using ultrasound. Prior to baseline measurements, mice were loosely restrained once a day over a period of 2 to 3 days to acclimate for handling. During this period, hair was removed from the chest using a depilatory agent. Echocardiograms were then performed on awake, loosely restrained mice prior to doxorubicin and treatment administration, then subsequently on days 3, 7, and 10 post-administration. Imaging was performed using a FujiFilm VisualSonics Vevo F2 equipped with an 18 mm linear probe with a bandwidth of 57-25 MHz. Two-dimensional, M mode parasternal short-axis views were recorded. Left ventricular dimensions, wall thickness, heart rate, and cardiac output were recorded. Left ventricle (LV) stroke volume (SV) was calculated as the difference between the end-diastolic volume (EDV) and the end-systolic volume (ESV). Ejection fraction (EF) was calculated as (EF = [(EDV-ESV)/EDV] *100).

### Acute Cardiotoxicity Studies

Doxorubicin was administered as a single intraperitoneal injection at a dose of 20 mg/kg. Immediately prior to the administration of doxorubicin, treatment was given via tail vein injection or oral gavage depending on the experimental group. Body weight was recorded prior to the study, and monitored three times per week throughout the experiment. At study completion, mice were sacrificed via carbon dioxide and cervical dislocation.

### Histology

Following euthanasia, hearts were dissected and phosphate buffered saline (PBS) was used to flush blood out of the heart. Hearts were fixed in neutral buffered formalin and paraffin embedded. Following embedding, 5 um thick cross sections were taken to visualize the left and right ventricles. Hematoxylin and eosin staining was performed to visualize cardiac muscle morphology. Images were taken at 1.6x magnification to show the entirety of the left and right ventricles, and 20x images were taken of the left ventricular free wall to visualize any structural abnormalities or show any presence of inflammation or vacuoles.

### Biodistribution Study Details

For the biodistribution study, NanoDMX liposomes were labeled using DiD dye at a 0.1% molar ratio relative to total lipids. Free DiD was administered as a control, and PBS was used as a negative control. Mice were injected with as single intraperitoneal injection of doxorubicin (20 mg/kg), followed immediately by intravenous administration of their respective experimental treatment. Animals were housed under normal conditions for 4 hours post-injection. Blood samples were collected 5 minutes after intravenous injections. Four hours later, mice were euthanized and dissected to remove the heart, liver, lungs, spleen, and kidneys, and fluorescence was measured using an IVIS Spectrum. Organs were weighed, and the percentage of total injected dose (%ID) per milligram of tissue was calculated based on fluorescence intensity, normalized to the total injected dose as determined from blood collected 5 minutes post-injection.

## Results

### Comparative Analysis of MAP4K4 Inhibitors in Reducing Doxorubicin-Induced Cell Death

To identify MAP4K4 inhibitors capable of mitigating doxorubicin-induced cardiotoxicity, we first assessed the cytotoxicity of four commercially available compounds in H9c2 cells by determining their LD_50_ values **(Fig. 1A)**. Among the inhibitors tested, DMX-5804 exhibited the highest LD₅₀ (46.51 µM), indicating the lowest intrinsic toxicity. Next, we evaluated whether these inhibitors could protect H9c2 cells from doxorubicin-induced cell death. H9c2 cells were co-treated with each MAP4K4 inhibitor at 10 µM and increasing concentrations of doxorubicin (10, 25, 50, and 75 µM) following the protocol as illustrated in **Fig. 1B**. As shown in **Fig. 1C**, DMX-5804 markedly preserved cell viability across all doxorubicin concentrations tested. In contrast GNE-495, MAP4K4-IN3, and PF-06260933 failed to improve cell viability compared to doxorubicin alone (**Figs. 1D-F**). Based on these findings, DMX-5804 was selected for subsequent mechanistic and functional studies.

**Figure 1:**
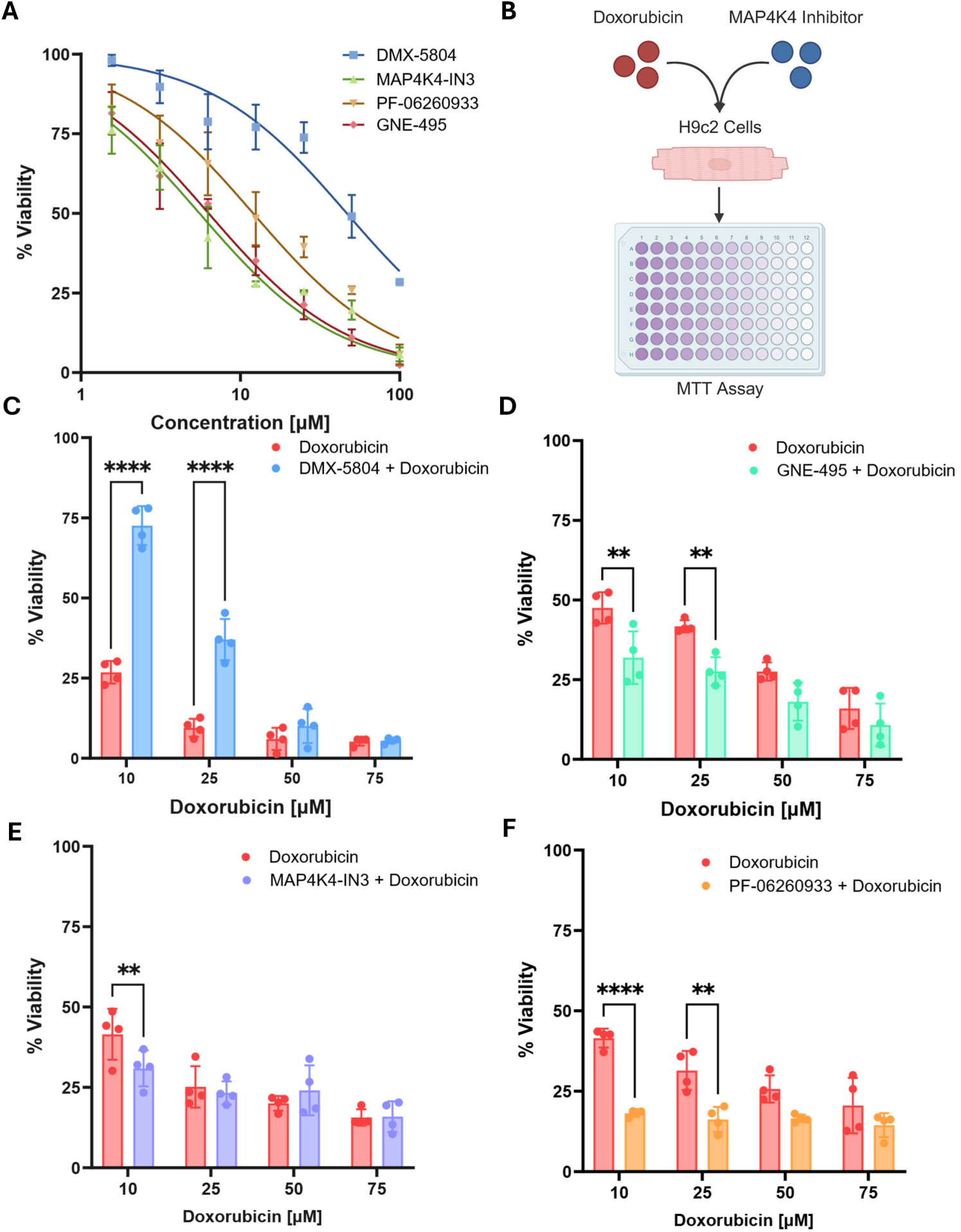
Comparative analysis of MAP4K4 inhibitors reveals differential effects on H9c2 cell viability in the presence of doxorubicin. **(A)** Effect of MAP4K4 Inhibitors on the viability of H9c2 Cells. **(B)** Schematic illustrating viability studies conducted on H9c2 cells treated with doxorubicin and MAP4K4 inhibitors. **(C)** Viability of H9c2 Cells Treated with Doxorubicin and DMX-5804. **(D)**Viability of H9c2 cells treated with doxorubicin and GNE-495, **(E)** Viability of H9c2 cells treated with doxorubicin and MAP4K4-IN3, **(F)** Viability of H9c2 cells treated with doxorubicin and PF-06260933. Statistical analysis was performed using two-way ANOVA to assess the effects of treatment and concentration, followed by Sidak’s multiple-comparisons post hoc test to compare groups at matched concentrations. Data are presented as mean ± stdev, **p<0.01, ***p<0.001, ****p<0.0001.

### Assessment of DMX-5804 in Preserving Mitochondrial Integrity Under Doxorubicin-Induced Stress

To evaluate the effect of DMX-5804 on mitochondrial integrity in cardiomyocytes exposed to doxorubicin, we assessed both mitochondrial mass and membrane potential. The following groups were tested: untreated control, 10 µM doxorubicin alone, 10 µM DMX-5804 alone, and a combination of 10 µM doxorubicin with 10 µM DMX-5804. Mitochondrial mass was visualized using MitoTrackerRed staining and nuclei were counterstained with DAPI (**Fig. 2A-D**). Representative images illustrate (A) untreated cells, (B) doxorubicin-treated cells, (C) DMX-5804-treated cells, and (D) co-treated cells. Quantification of mean MitoTracker Red fluorescence across conditions is shown in **Fig. 2E**.

**Figure 2:**
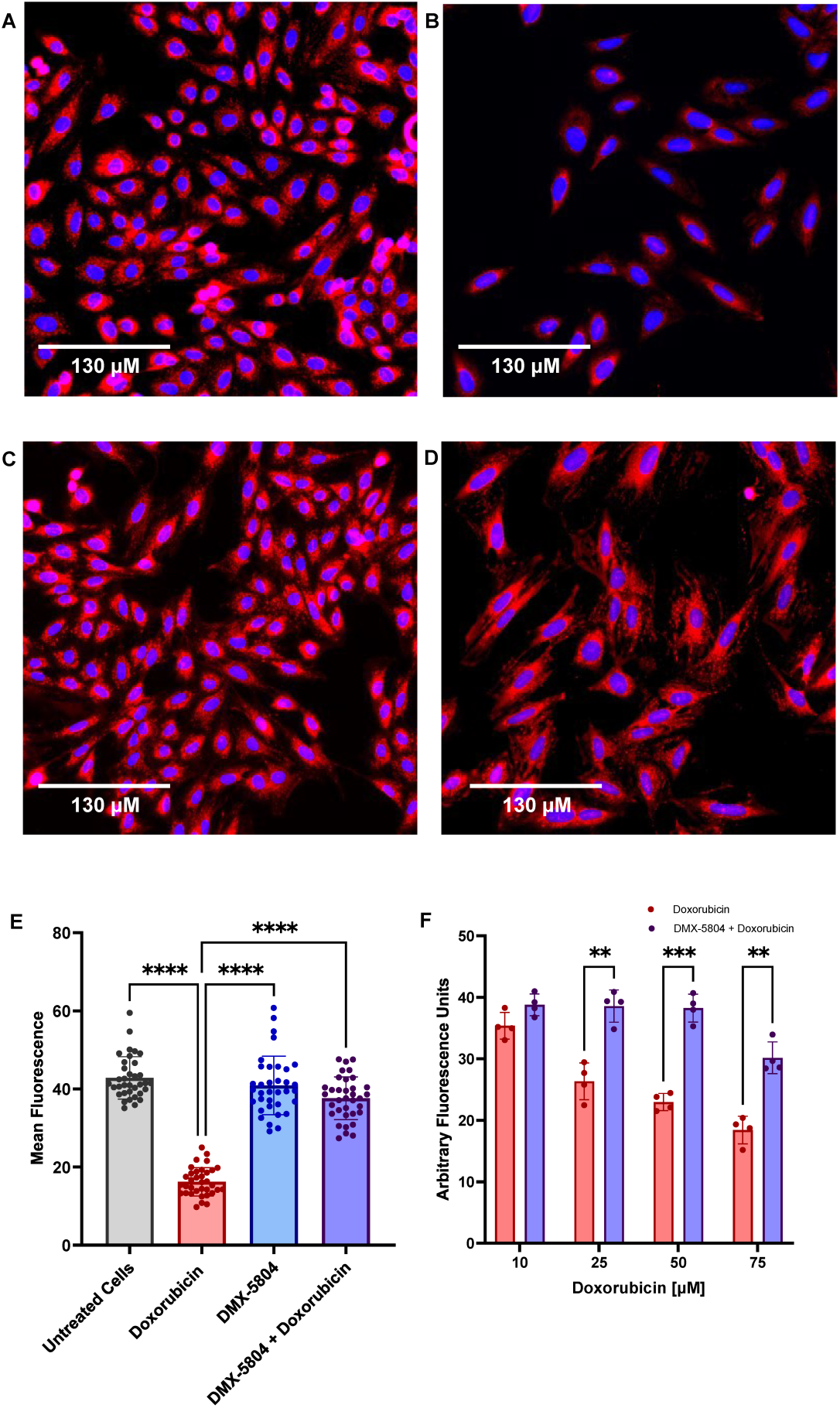
Effects of DMX-5804 on mitochondrial health of H9c2 cells that have been exposed to doxorubicin. (A) MitoTrackerRed and DAPI staining of healthy, control H9c2 cells. (B) MitoTrackerRed and DAPI staining of H9c2 cells exposed to 10 µM doxorubicin for 24 hours. (C) MitoTrackerRed and DAPI staining of H9c2 cells exposed to 10 µM DMX-5804 for 24 hours. (D) MitoTrackerRed and DAPI staining of H9c2 cells exposed to 10 µM doxorubicin and 10 µM DMX-5804 for 24 hours. (E) Quantification of MitoTrackerRed staining across all four conditions of drug treatment. Data are presented as mean ± stdev, **p<0.01, ***p<0.001, ****p<0.0001 with one-way ANOVA followed by Dunnett’s multiple comparison’s post-test. (F) TMRM assay of cells exposed to increasing concentrations of doxorubicin alone or increasing concentrations of doxorubicin and 10 µM DMX-5804. Data are presented as mean ± stdev, **p<0.01, ***p<0.001, ****p<0.0001 with two-way ANOVA followed by Sidak’s multiple comparison’s post-test.

To further assess mitochondrial function, we performed a TMRM assay using 10 µM DMX-5804 combined with increasing concentrations of doxorubicin **(Fig. 2F)**. As doxorubicin concentrations increased, mitochondrial membrane potential progressively declined; however co-treatment with DMX-5804 significantly preserved mitochondrial membrane potential compared to doxorubicin alone. Collectively these findings indicate that DMX-5804 mitigates doxorubicin-induced mitochondrial damage by maintaining both mitochondrial mass and membrane potential in cardiomyocytes.

### Effects of DMX-5804 on MAP4K4 Pathway Activation and Doxorubicin Accumulation

To assess whether DMX-5804 protects doxorubicin-exposed H9c2 cells through the JNK pathway suppression via MAP4K4 inhibition, we performed western blotting and RT-qPCR analyses. Lysates from H9c2 cells treated with doxorubicin alone, DMX-5804 alone, or both compounds for 4 and 24 hours were analyzed by western blot **(Figs. 3E)**, and gene expression related to the JNK pathway, mitochondrial mitophagy, and solute carriers was quantified using RTqPCR.

**Figure 3:**
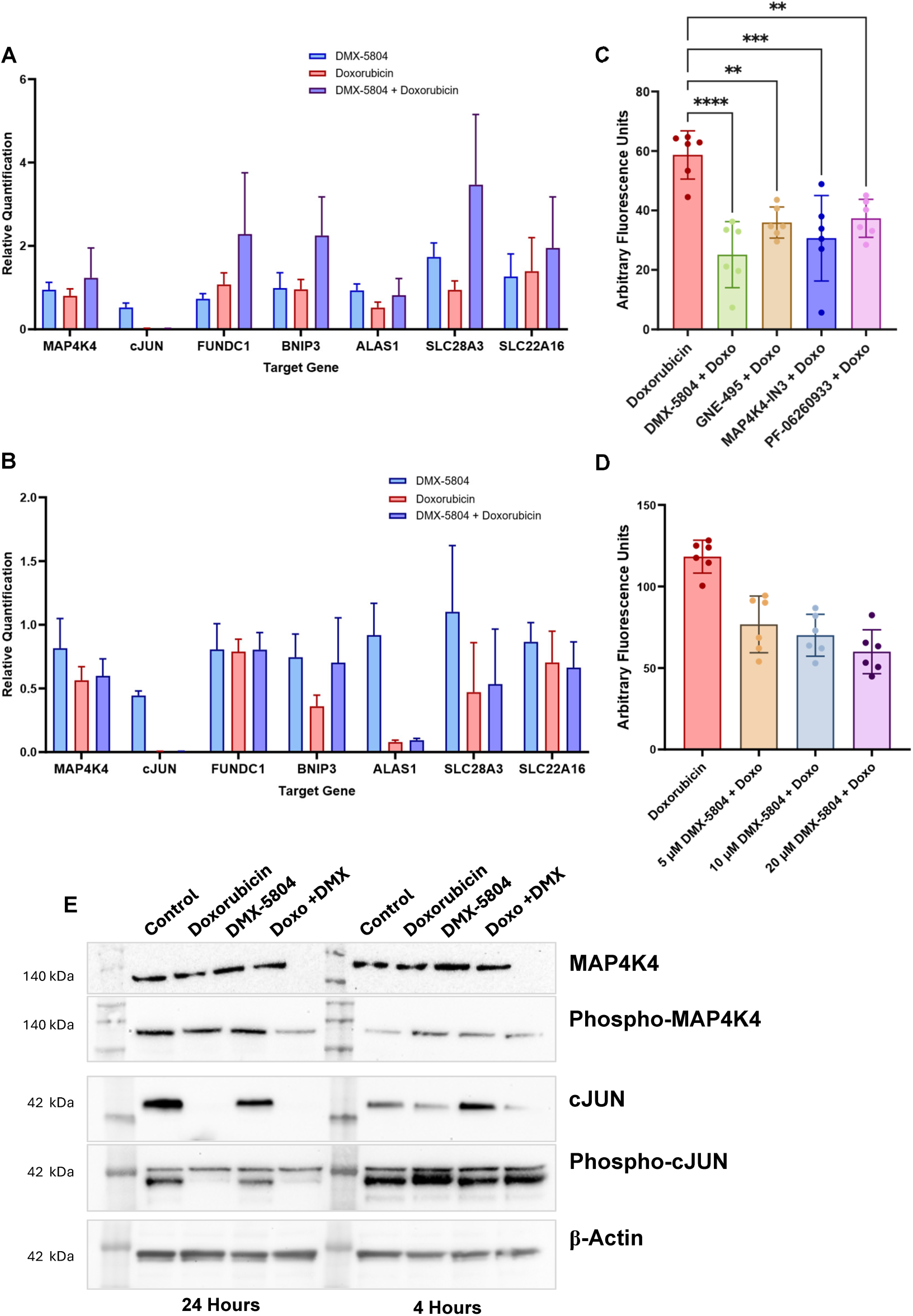
Effects of DMX-5804 on doxorubicin-induced gene expression, drug uptake, and protein signaling in H9c2 cardiomyoblasts. **(A)** qPCR analysis of H9c2 cells treated with doxorubicin, DMX-5804, or both for 4 hours. **(B)** qPCR analysis of H9c2 cells treated with doxorubicin, DMX-5804, or both for 24 hours. RT-qPCR data are shown as mean ± SD from n = 3 independent biological replicates, each measured in duplicate technical replicates. **(C)** Uptake of 10 μM doxorubicin into H9c2 cells co-treated with 10 μM MAP4K4 inhibitors (DMX-5804, GNE-495, MAP4K4-IN3, and PF-06260933) after 24 hours. Data represent mean ± standard deviation from six technical replicates per condition from a single experiment; statistical comparisons were performed using one-way ANOVA. **(D)** Uptake of 10 μM doxorubicin into H9c2 cells co treated with increasing concentrations of DMX-5804 (5 μM, 10 μM, and 20 μM) after 24 hours. Data represent mean ± standard deviation from six technical replicates per condition from a single experiment; statistical comparisons were performed using one-way ANOVA followed by Dunnett’s multiple comparison’s post-test. **(E)** Western blot analysis of H9c2 cells treated with doxorubicin, DMX-5804, or both for 24 hours and 4 hours. Membranes were probed for β-actin as a loading control to verify equal protein loading across lanes. Blots shown are representative of at least three independent experiments.

Western blots revealed that doxorubicin markedly suppressed total cJUN expression, with partial suppression at 4 hours and near-complete suppression at the 24 hours. Phosphorylated cJUN was initially elevated at 4 hours but decreased by 24 hours. RTqPCR results mirrored these findings, showing strong downregulation of cJUN transcripts in doxorubicin and doxorubicin + DMX-5804 groups at both time points.

Interestingly, mitophagy-related genes FUNDC1 and BNIP3 exhibited an early increase in the doxorubicin + DMX-5804 group **(Fig. 3A)**, returning to baseline by the 24 hours **(Fig. 3B)**. Similarly, solute carrier genes SLC28A3 and SLC22A16 showed transient upregulation at 4 hours in co-treated cells, normalized by 24 hours.

Doxorubicin uptake studies (**Figs. 3C-D**) demonstrated reduced intracellular doxorubicin in H9c2 cells co-treated with MAP4K4 inhibitors, including DMX-5804 with modest dose-dependent decreases observed. Collectively these data indicate that DMX-5804-mediated cardioprotection is not driven by JNK pathway as previously hypothesized but is instead linked to mitochondrial protection and altered solute carrier expression, leading to reduced intracellular doxorubicin accumulation.

### Effects of DMX-5804 and Other MAP4K4 Inhibitors on Viability and Drug Accumulation in Cancer Cells

To determine whether DMX-5804 is cardioprotective, without conferring protection to cancer cells, we repeated experiments previously performed in H9c2 cells using triple negative breast cancer cell lines. **Fig. 4A**, shows the testing of MAP4K4 inhibitors in MDA-MB-231 cells to determine LD_50_ values. DMX-5804 exhibited an LD_50_ value of 46.51 µM in H9c2 cells, whereas in MDA-MB-231 cells the values were markedly lower at 11.59 µM. **Fig. 4B** illustrates the uptake of doxorubicin in MDA-MB-231 cells treated with doxorubicin alone or in combination with MAP4K4 inhibitors. Unlike H9c2 cells, where MAP4K4 inhibitors reduced doxorubicin uptake, no such effect was observed in MDA-MB-231 cells.

**Figure 4:**
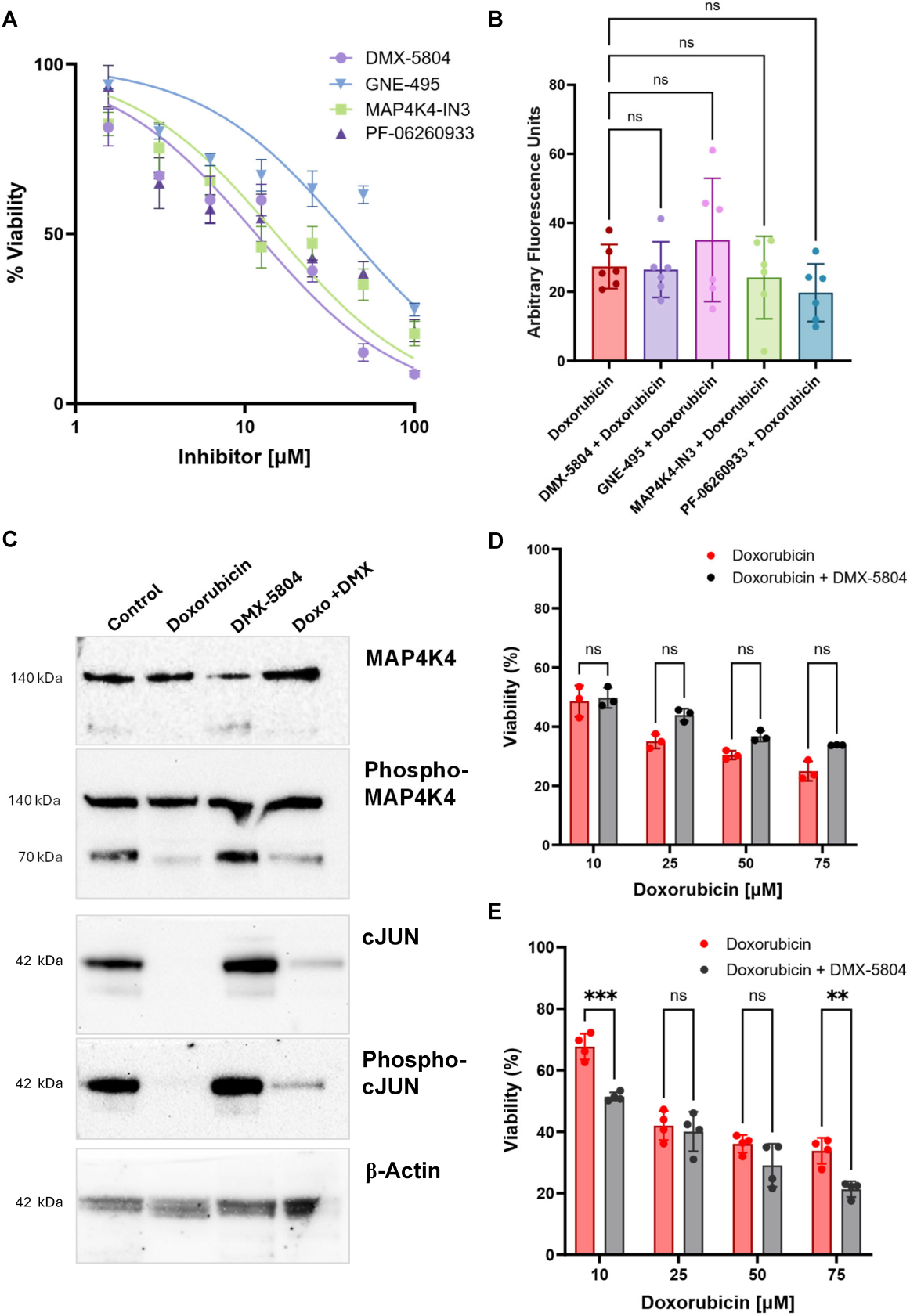
Effects of MAP4K4 inhibition on doxorubicin uptake, cell viability, and protein signaling in breast cancer cells. **(A)** Effect of MAP4K4 inhibitors on the cell viability of MDA-MB-231 cells. **(B)** Uptake of doxorubicin into MDA-MB-231 cells co-treated with MAP4K4 inhibitors after 24 hours. **(C)** Viability of MDA-MB-231 cells treated with doxorubicin and DMX-5804. **(D)** Viability of 4T1 cells treated with Doxorubicin and DMX-5804. **(E)** Western blot analysis of MDA-MB-231 cells treated with doxorubicin, DMX-5804, or both for 24 hours. Data are presented as mean ± stdev, n=4. Statistical analysis was performed using two-way ANOVA to assess the effects of treatment and concentration, followed by Sidak’s multiple-comparisons post hoc test to compare groups at matched concentrations. Data are presented as mean ± stdev, **p<0.01, ***p<0.001, ****p<0.0001.

**Figures 4D** and **4E** present viability assays performed with doxorubicin alone and doxorubicin plus DMX-5804 in MDA-MB-231 and 4T1 cells, respectively. In both cases, DMX-5804 did not protect against doxorubicin-induced cell death. **Figure 4C** shows western blot analysis of MDA-MB-231 cells after 24 hours of treatment with doxorubicin, DMX-5804, or their combination. Similar to H9c2 cells, treatment resulted in a significant reduction in cJUN and phosphorylated cJUN protein levels. Overall, we demonstrate that while DMX-5804 is cardioprotective, it does not preserve the viability of cancer cells exposed to doxorubicin.

### Physicochemical Characterization of Liposomal Formulations Loaded with DMX-5804

After confirming that DMX-5804 provides cardioprotection in the presence of doxorubicin, we aimed to develop a formulation suitable for intravenous (IV) administration and co-dosing with doxorubicin. This need arose because DMX-5804 is highly hydrophobic - readily soluble in organic solvents such as DMSO (up to ∼100 mg/mL) but insoluble in water - making direct IV delivery impractical. Clinically, this is critical because doxorubicin is administered intravenously, and cardiotoxicity is a major dose-limiting side effect that compromises cancer treatment outcomes. A formulation that enables simultaneous IV co-administration of DMX-5804 with doxorubicin could allow real-time cardioprotection during chemotherapy without altering standard treatment protocols.

To address this, we encapsulated DMX-5804 into liposomes using thin-film hydration followed by sonication and extrusion, enabling aqueous dispersion and systemic administration. As DMX-5804 is a hydrophobic drug, it was solubilized in an organic solvent, mixed with the lipids, and dried to form a drug–lipid thin film. The film was then rehydrated in an aqueous buffer, followed by sonication and extrusion to produce uniformly sized liposomes as previously described.^17^ Following encapsulation, multiple liposomal formulations were generated at varying drug-to-lipid weight ratios, and their size, polydispersity index (PDI), and zeta potential were characterized. **Figure 5A** displays size (left axis) and PDI (right axis) for these formulations. Notably, formulations containing higher amounts of phosphatidylserine exhibited increased PDI at higher drug:lipid weight ratios. Since our primary objective was to maximize DMX-5804 loading, we selected a formulation with reduced phosphatidylserine content: DSPC:DSPS:Cholesterol at a 60:20:20 molar ratio and a 50% drug-to-lipid weight ratio for downstream studies. This optimized formulation was designated “NanoDMX” for clarity in subsequent experiments.

**Figure 5:**
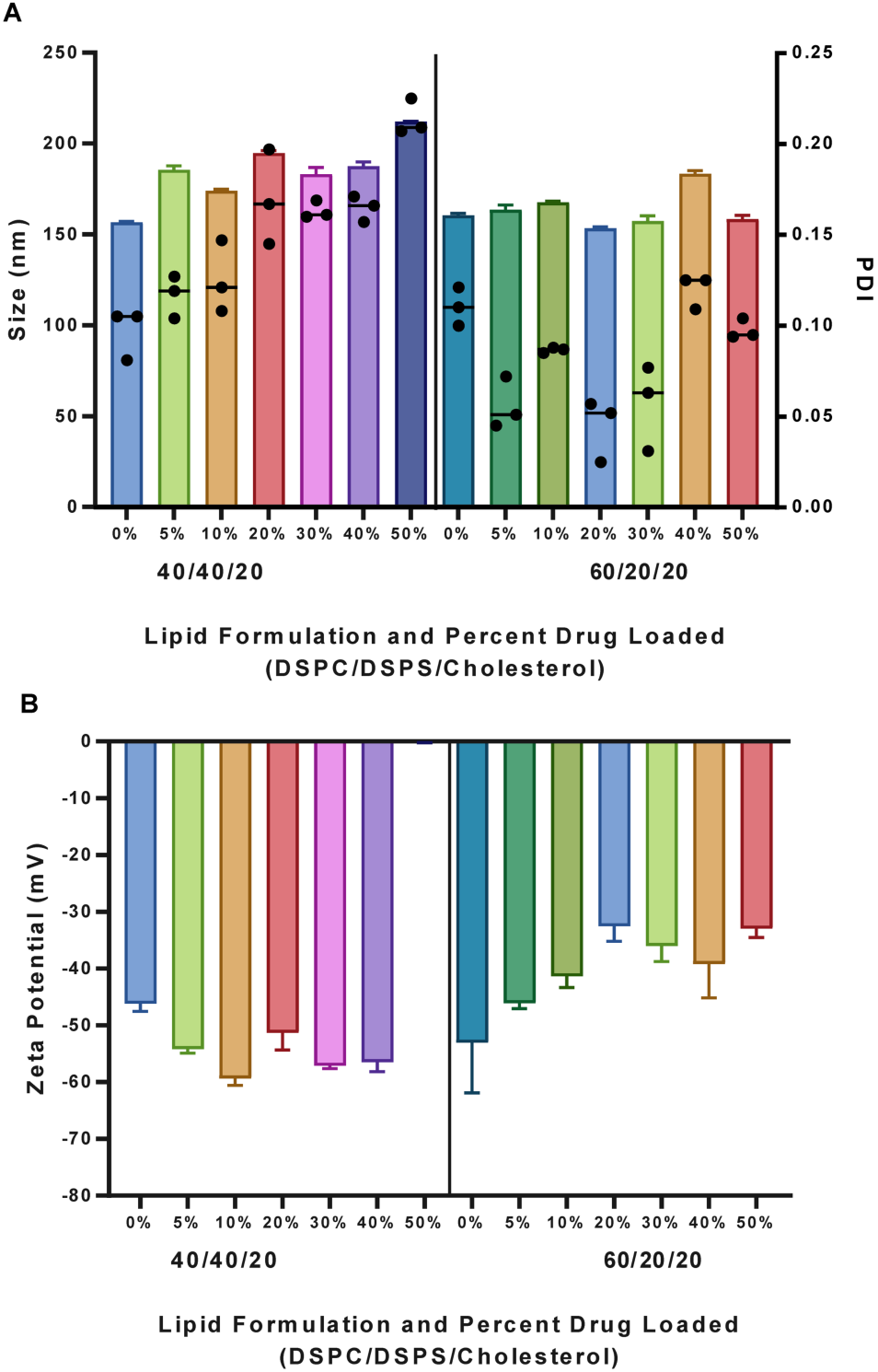
Physicochemical characterization of DMX-5804–loaded liposomal formulations developed for intravenous co-administration with doxorubicin. **(A)** Size and PDI of liposomal formulations tested. (B) Zeta potential of liposomal formulations. Liposome size, polydispersity index (PDI), and zeta potential are reported as mean ± standard deviation from n = 3 independent liposome preparations.

The primary objective of the initial in vivo study was to determine whether the modest inclusion of phosphatidylserine in the NanoDMX formulation could leverage its reported anti-inflammatory properties. To address this, we included a phosphatidylserine free NanoDMX formulation and its vehicle, consisting of a 70:30 DSPC:Cholesterol ratio, while maintaining the same drug-to-lipid ratio as NanoDMX. An acute, single high-dose doxorubicin toxicity model was used, and mice were monitored over the course of ten days. The primary endpoints were left ventricular ejection fraction (EF%) and fractional shortening (FS%), clinically relevant indicators of cardiac function. Briefly, mice received an intraperitoneal injection of doxorubicin (20 mg/kg), followed by intravenous administration of their assigned treatment. Liposome-containing groups were dosed at 25 mg/kg based on lipid content. **Figure 6A** illustrates ejection fraction (EF%) over the course of ten days. By day 3, EF% sharply declined in the PBS and Alternate NanoDMX vehicle groups, indicating declining functional output of the heart.

**Figure 6:**
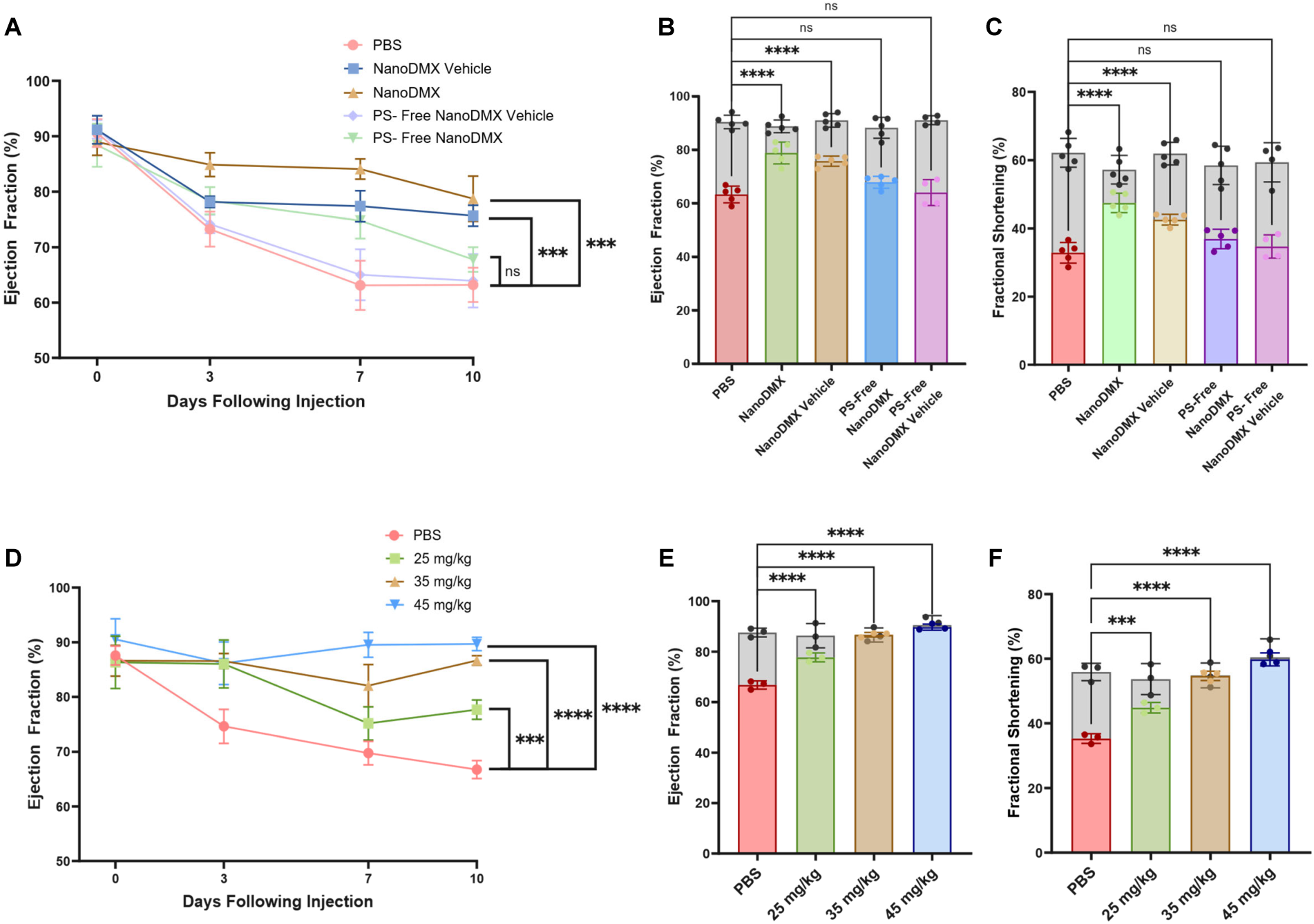
Effects of NanoDMX on doxorubicin-induced cardiac dysfunction in mice. (A) Ejection fraction of mice treated with doxorubicin, doxorubicin and NanoDMX, doxorubicin and phosphatidylserine free NanoDMX, and relevant vehicle controls. (B) Comparison of ejection fraction of mice at day 0 and day 10, with grey bars representing day 0 and colored bars representing day 10. (C) Comparison of fractional shortening of mice at day 0 and day 10. Data are presented as mean ± standard deviation from n = 5 mice per group and were analyzed using one-way ANOVA followed by Sidak’s multiple-comparisons post hoc test. (D) Ejection fraction over a ten-day period in mice treated with doxorubicin and doxorubicin combined with NanoDMX at increasing doses. (E) Ejection fraction of mice treated with increasing doses of NanoDMX and doxorubicin at day 0 and day 10. (F) Fractional shortening of mice treated with doxorubicin and increasing doses of NanoDMX at day 0 and day 10. Data are presented as mean ± standard deviation from n = 3 mice per group and were analyzed using one-way ANOVA followed by Sidak’s multiple-comparisons post hoc test.

In contrast, the NanoDMX, NanoDMX vehicle, and Alternate NanoDMX group maintained a preservation of ejection fraction (EF%) until day 10, when the Alternate NanoDMX group also began to decline. **Figure 6B** shows the comparison of ejection fraction on day 0 and day 10, showing that only NanoDMX and NanoDMX vehicle group show a statistically significant improvement in ejection fraction when compared to the doxorubicin + PBS group, with the NanoDMX demonstrating the strongest effect. Similarly, **Figure 6C** presents fractional shortening (FS%) changes across groups. Collectively, these data suggest that phosphatidylserine inclusion in NanoDMX contributes substantially to the observed cardioprotective efficacy following high-dose doxorubicin exposure, with further improvements mediated by DMX508.

We next investigated whether NanoDMX exhibits a dose-dependent protective effects against doxorubicin-induced cardiotoxicity when co-administered with doxorubicin. Using the same model as in the previous study, mice received a 20 mg/kg intraperitoneal injection of doxorubicin followed by an intravenous injection of either PBS or NanoDMX liposomes at varying doses. For this study, we tested the previously used dose of 25 mg/kg along with 35 and 45 mg/kg, calculated based on liposome amount. Representative echocardiography images at day 10 for each group (PBS, 25 mg/kg, 35 mg/kg, and 45 mg/kg) are shown in **Figure 6 D-E**. **Fig. 6D** illustrates the progression of ejection fraction over 10 days, measured throughout the study. As expected, the doxorubicin + PBS group exhibited a sharp decline in ejection fraction after day 3. In contrast, all NanoDMX-treated groups preserved ejection fraction, with a clear dose-dependent effect: mice receiving 45 mg/kg of NanoDMX maintained near-normal function throughout the study.

Comparisons of ejection fraction and fractional shortening at days 0 and 10 are shown in **Figure 6E**, confirming that NanoDMX treatment preserved these clinically relevant functional metrics. Additionally, **Figure 6F** shows body weight changes over the 10-day period, where NanoDMX-treated groups demonstrated better weight maintenance compared to PBS controls. Overall, these findings indicate a dose-dependent benefit of NanoDMX in preserving cardiovascular function and body weight in doxorubicin-exposed mice.

### Impact of NanoDMX on Ejection Fraction and Fractional Shortening in a Cardiotoxicity Model

After identifying the best formulation and dose, we evaluated the prophylactic efficacy of DMX-5804 against doxorubicin-induced cardiotoxicity when administered with and without a liposomal carrier. Because DMX-5804 is hydrophobic and unsuitable for intravenous delivery without a carrier, it was administered orally at 22 mg/kg - the same effective dose present in NanoDMX, which was formulated at a 50% drug-to-lipid ratio and previously optimized at 45 mg/kg based on liposomal quantities. Both treatments followed the same timeline: intraperitoneal doxorubicin (20 mg/kg) was given, immediately followed by the assigned therapeutic. **Fig. 7A** shows the ten-day progression of ejection fraction across 4 time points. Consistent with prior findings, the doxorubicin + PBS group exhibited a marked decline in ejection fraction after day 3. Oral DMX-5804 mirrored this decline, showing minimal cardioprotection compared to NanoDMX and its empty vehicle. **Fig. 7B and C** summarize ejection fraction and fractional shortening at baseline (Day 0) and Day 10. **Fig. 7D** shows body weight changes, indicating that oral DMX-5804 did not mitigate doxorubicin-associated weight loss. Despite DMX-5804’s in vitro cardio-protective effects, un-encapsulated DMX-5804 failed to confer benefit in vivo. In contrast, across three independent studies, the inclusion of DMX-5804 in phosphatidylserine liposomes enhanced DMX-5804’s ability to protect against doxorubicin cardiotoxicity.

**Figure 7:**
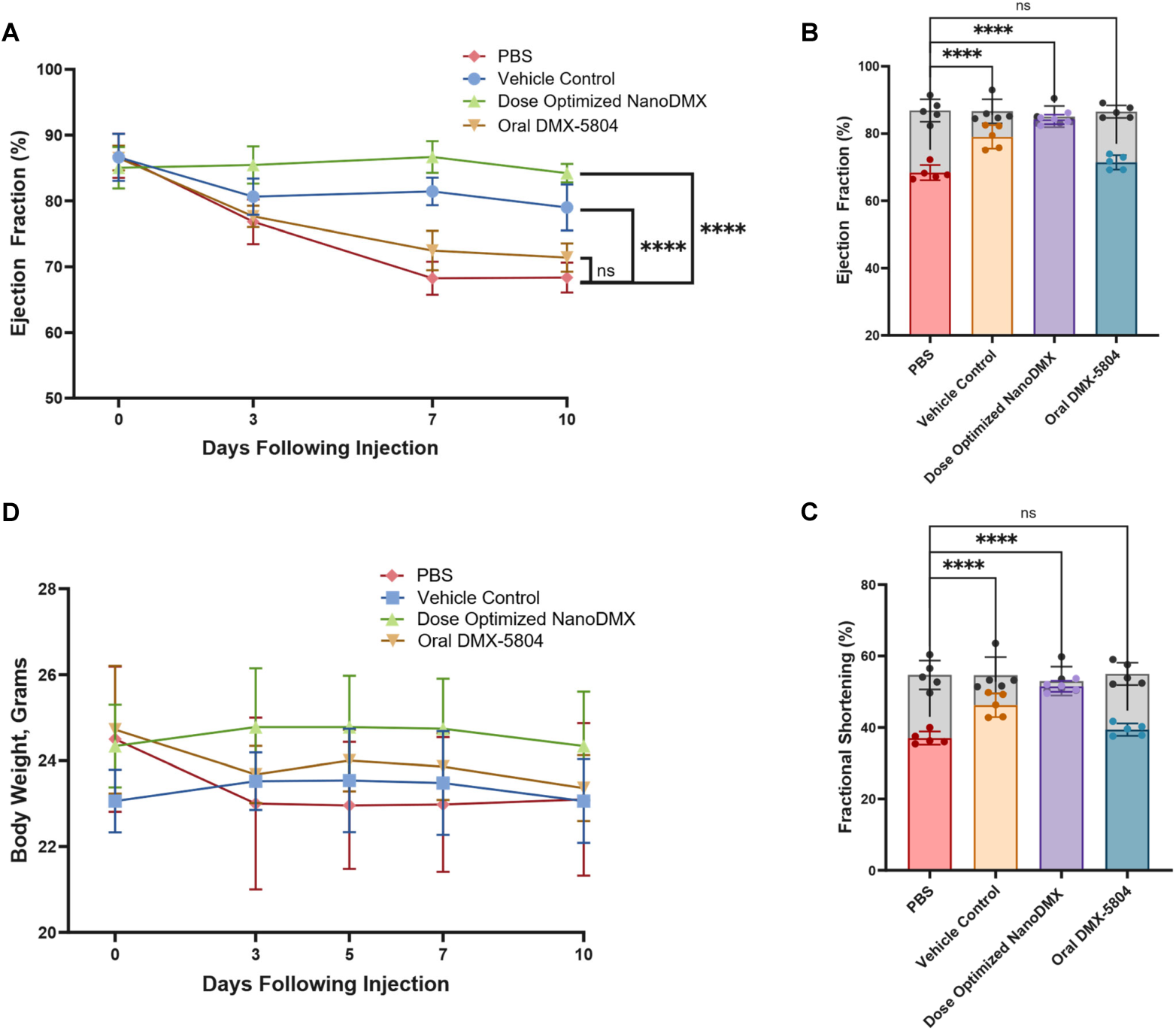
Effects of DMX-5804 delivery on doxorubicin-induced changes in cardiac function and body weight over a 10-day period. (A) Ejection fraction over the 10-day period. (B) Comparison of ejection fraction at baseline (Day 0) and Day 10 across all groups. (C) Comparison of fractional shortening at baseline (Day 0, grey bars) and Day 10 (colored bars) across all groups. (D) Body weight of mice over the ten-day evaluation period following treatment. Data are presented as mean ± standard deviation from n = 5 mice per group and were analyzed using one-way ANOVA followed by Sidak’s multiple-comparisons post hoc test.

### Impact of NanoDMX on Cardiac Tissue Morphology in a Cardiotoxicity Model

**Figure 8** illustrates changes in vivo in cardiac morphology following doxorubicin administration, as well as the preservation of cardiac structure when doxorubicin is co-administered with NanoDMX. Healthy, normal morphology of cardiac muscles is shown in **Figure 8A** depicts healthy cardiac tissue with normal morphology, characterized by well-organized and uniformly aligned cardiomyocytes. Conversely, **Figure 8B** shows heart from doxorubicin-treated mice, which exhibit marked disruption of cardiac muscle architecture, along with the presence of vacuolization and inflammatory infiltrates, as denoted by the white and black arrows, respectively. **Figure 8C** demonstrates a similar pattern mice treated with doxorubicin in combination with orally administered, unmodified DMX-5804, where prominent vacuoles and muscle damage are evident as clear, unstained regions. In **Figure 8D**, however, there is an observed preservation of cardiac muscle fiber organization and cardiomyocyte architecture, as well as minimal vacuolization. Overall, these findings indicate that the co-treatment with encapsulated DMX-5804 (NanoDMX) effectively preserves cardiac tissue morphology following doxorubicin exposure, whereas orally administered DMX-5804 does not confer comparable structural protection.

**Figure 8:**
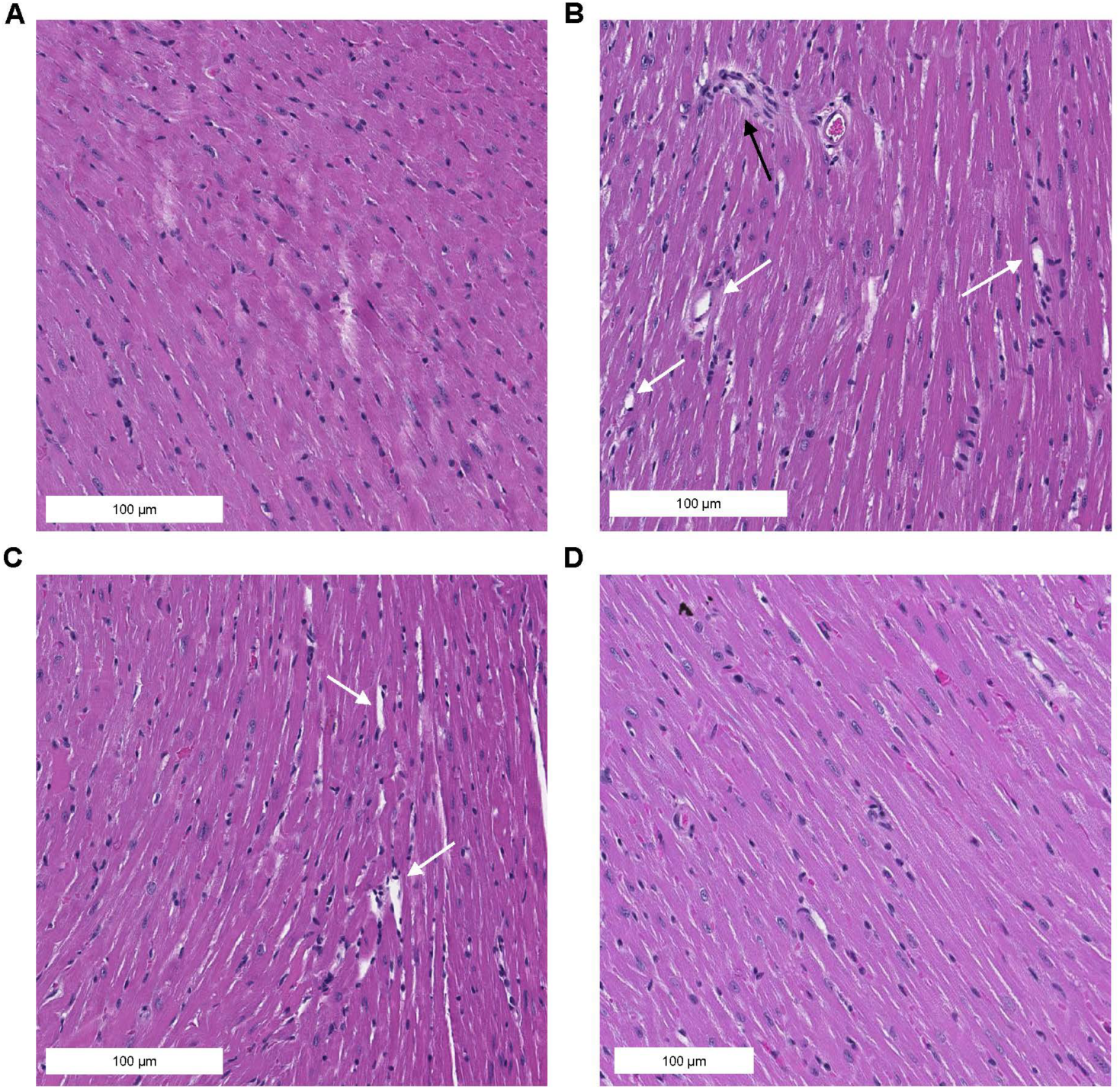
Representative H&E-stained sections of the left ventricular free wall from mice receiving doxorubicin with or without DMX-5804 treatment. (A) 20x magnification of left ventricular free wall of healthy mice. (B) 20x magnification of left ventricular free wall of doxorubicin mice. (C) 20x magnification of left ventricular free wall of doxorubicin and orally administered DMX-5804 mice. (D) 20x magnification of left ventricular free wall of doxorubicin and NanoDMX treated mice.

## DISCUSSION

Cardiovascular drug development has historically progressed more slowly than advances in other therapeutic areas, particularly in the context of cardiotoxicity and myocardial injury.^18–21^ This gap underscores the need for mechanistically informed strategies that directly address the unique vulnerabilities of cardiac tissue.

Here, we have developed an intravenously deliverable, dual-action prophylactic formulation against doxorubicin-induced cardiotoxicity by encapsulating the MAP4K4 inhibitor DMX-5804 in phosphatidylserine-containing liposomes (NanoDMX). This approach overcame the solubility limitations of DMX-5804 and enabled effective IV administration. From the MAP4K4 inhibitors tested only DMX-5804 preserved cardiomyocyte viability under doxorubicin exposure. Through our in vitro work, we found that the use of DMX-5804 in cardiomyocytes exposed to doxorubicin led to a preservation in viability, unexpected changes in the expression of the transcription factor cJUN and its associated activation, as well as changes in the expression of mitophagy regulation proteins and solute carrier proteins, and protection against mitochondrial damage in cardiac cells.

In contrast, DMX-5804 did not protect triple-negative breast cancer cells from doxorubicin-induced damage, supporting its potential as a cardio-oncology therapeutic. Our in vivo studies revealed that the combined use of DMX-5804 and phosphatidylserine containing liposomes is able to protect mice from doxorubicin induced cardiovascular damage, as evidenced by the preservation of left ventricular ejection fraction and fractional shortening. The combination of this in vitro and in vivo work shows promise in the use of an intravenous prophylactic to protect against anthracycline induced cardiotoxicity while not also protecting cancerous cells.

Our initial in vitro evaluation of 4 commercially available MAP4K4 inhibitors **(Fig. 1A-F)** revealed that only DMX-5804 effectively preserved cardiac cell viability and mitochondrial function. The other inhibitors failed to confer protection, which may reflect differences in target specificity. This suggests that these compounds are not exclusively inhibiting MAP4K4 and may engage off-target pathways, contributing to either the absence of variability of protective effects. One possible mechanism involves interaction between these kinase inhibitors and surface solute carrier proteins. This hypothesis is supported by **Fig. 3C**, where co-treatment with MAP4K4 inhibitors reduced intracellular accumulation of doxorubicin in H9c2 cells, indicating potential modulation of drug uptake.

DMX-5804 was selected as the primary candidate for downstream applications because it demonstrated the greatest ability to preserve cardiac cell viability among the tested compounds. To investigate whether its protective effect was mediated through mitochondrial preservation, we performed mitochondrial staining and membrane potential assays. Co-treatment of H9c2 cells with DMX-5804 and doxorubicin maintained both mitochondrial number and membrane potential after 24 hours of exposure. These findings indicate that, although additional signaling pathways beyond the predicted JNK pathway may contribute, DMX-5804 protects cardiac cells at least in part by preserving mitochondrial integrity. Mitochondrial preservation is critical for cardiomyocytes, as these cells have one of the highest mitochondrial densities of any cell type, with mitochondria comprising approximately 30 to 40% of their total volume.^22^

Interestingly, DMX-5804 treatment of cardiomyocytes in vitro did not lead to a reduction in cJUN expression in cardiac cells, whereas doxorubicin markedly reduced it. MAP4K4, and the MAPK family as a whole, has been attributed to the regulation of the JNK pathway. Independently, MAP4K4 specifically has been shown in different tissue types to directly regulate the activation and subsequent phosphorylation of cJUN in response to different types of cellular stress.^23^ There is research to suggest that doxorubicin would lead to an increase in cJUN expression, as it is a known activator of the JNK pathway and cJUN phosphorylation.^24^ This unexpected finding highlights the complex interplay between cJUN, the JNK pathway as a whole, mitochondrial-driven cell death, and doxorubicin - mechanisms that warrant further investigation. At 4 hours, doxorubicin, whether administered alone or with DMX-5804, began to suppress protein levels of cJUN, and by 24 hours, cJUN protein levels were suppressed almost entirely.

Notably, DMX-5804 treatment reduced doxorubicin accumulation in H9c2 cells. While similar effects have been reported for other kinase inhibitors, often attributed to inhibition of surface uptake transporters such as SLC proteins^25^, our data indicate that this reduction is not due to SLC downregulation. In fact, RT-qPCR analysis revealed upregulation of SLC28A3 and SLC22A16 under combined doxorubicin and DMX-5804 treatment. This suggests that these transporters may be preserved to support essential functions, such as nucleoside and carnitine uptake, which are critical for mitochondrial fatty acid transport and energy production in cardiomyocytes.^26^ Thus, while decreased doxorubicin uptake is cardioprotective, the mechanism appears independent of SLC suppression and may involve alternative pathways.

A crucial component of an effective cardioprotective therapeutic for cancer patients is that it does not confer protection to cancer cells. To evaluate the feasibility of using DMX-5804 prophylactically, we evaluated DMX-5804 in MDA-MB-231 cells (**Fig. 4A-F**). Strikingly, DMX-5804 exhibited a much lower LD_50_ in MDA-MB-231 cells (11.59 µM) than in H9c2 cells (46.51 µM), and it failed to preserve cancer cell viability when co-treated with doxorubicin. This demonstrates that inherently, DMX-5804 is more toxic to cancerous cells than to cardiomyocytes. Interestingly, we also found across all four of our initially tested inhibitors that the co-treatment of MAP4K4 inhibitor with doxorubicin did not reduce the accumulation of doxorubicin intracellularly in MDA-MB-231 cells. This suggests that the uptake pathway being blocked by a MAP4K4 inhibitor in cardiomyocytes is either not absent or significantly less active in MDA-MB-231 cells, reinforcing the selectivity of this approach. This highlights additional potential mechanisms that could be investigated in future studies to further understand the mechanisms by which doxorubicin kills cardiomyocytes. Western blot analysis revealed a similar doxorubicin pattern to that observed in cardiomyocytes, with cJUN and phosphorylated cJUN protein levels markedly suppressed by 24 hours in all groups exposed to doxorubicin. In contrast, DMX-5804 alone or in combination with doxorubicin did not reduce levels of cJUN or phosphorylated cJUN levels, consistent with findings in cardiomyocytes.

This eventual downregulation of cJUN has been observed previously and prior research indicates that cJUN plays a critical role in preventing stress-induced cardiac remodeling.^27^ While MAP4K4 inhibition was expected to suppress cJUN as part of a protective mechanism for cardiomyocytes, doxorubicin-induced suppression may represent a maladaptive response that contributes to improper remodeling. Complete deletion of cJUN is embryonically lethal, as the protein is essential for the formation of beating cardiomyocytes during development.^28^ Therefore, to effectively leverage MAP4K4 inhibition as a therapeutic strategy, a deeper understanding of how to modulate cJUN expression at different stages of cardiomyocyte stress is needed.

In developing a liposomal formulation for DMX-5804 delivery, our goal was to design a carrier with inherent therapeutic benefits. To achieve this, we incorporated phosphatidylserine into the liposomes. Phosphatidylserine is a well-characterized anti-inflammatory lipid and has demonstrated efficacy in preclinical models of myocardial infarction.^29^ Additionally, prior studies have shown that phosphatidylserine-containing liposomes can modulate cardiac macrophage activity, helping to resolve maladaptive early inflammatory responses to cardiomyocyte distress.^30^ ^31^ While our study did not directly assess macrophage-mediated effects, this mechanism is supported by extensive literature. Importantly, our comparison of NanoDMX with phosphatidylserine-free liposomes revealed that phosphatidylserine inclusion significantly enhanced NanoDMX’s efficacy against doxorubicin-induced cardiotoxicity.

Notably, NanoDMX significantly outperformed orally administered DMX-5804 (Fig. 8). DMX-5804-treated animals exhibited similar drops in body weight and cardiovascular function as mice that received doxorubicin and PBS alone. These findings highlight that intravenous delivery via a liposomal carrier provides DMX-5804 with a renewed potential as a therapeutic strategy for the prevention of anthracycline-induced cardiotoxicity.

## CONCLUSION

In conclusion, our data demonstrate that the MAP4K4 inhibitor DMX-5804 protects cardiomyocytes from doxorubicin-induced cell death, primarily through mitochondrial preservation rather than inhibition of cJUN activation. DMX-5804 also reduced intracellular doxorubicin accumulation in cardiomyocytes without affecting uptake in triple-negative breast cancer cells. Upregulation of mitophagy-related proteins FUNDC1 and BNIP3 further supports a mitochondrial-driven protective mechanism independent of the JNK pathway.

In vivo, NanoDMX, a liposomal formulation combining DMX-5804 with phosphatidylserine, maintained cardiac function following high-dose doxorubicin exposure, with efficacy increasing in a dose-dependent manner. Importantly, NanoDMX outperformed orally administered DMX-5804, underscoring the advantage of intravenous liposomal delivery for preventing anthracycline-induced cardiotoxicity.

## DATA AVAILABILITY

All raw data and images for this article are available at https://doi.org/10.15139/S3/FQKK2K.

## ACKNOWLEDGEMENTS

JN and BCJ acknowledge funding from the NHLBI through awards R01HL161456 and R01HL165294. JTK received funding from the UNC Dissertation Completion Fellowship and through the T32 fellowship funded from NIGMS T32GM122741. Figure 1B was created through Biorender. Ultrasound Imaging is supported by the Preclinical Molecular Imaging Core Facility at UNC Biomedical Imaging Research Center (BRIC), and the ultrasound imaging system was funded by an NIH shared instrumentation grant, S10OD034328. Histopathology/Digital Pathology was performed by the Pathology Services Core at the University of North Carolina-Chapel Hill, which is supported in part by an NCI Center Core Support Grant (P30CA016086)

## References

1. Tetterton-Kellner J, Jensen BC, Nguyen J. Navigating cancer therapy induced cardiotoxicity: From pathophysiology to treatment innovations. Adv Drug Deliv Rev. 2024;211:115361. doi:10.1016/j.addr.2024.115361

2. Zaorsky NG, Churilla TM, Egleston BL, et al. Causes of death among cancer patients. Ann Oncol. 2017;28(2):400–407. doi:10.1093/annonc/mdw604

3. Eneh C, Lekkala MR. Dexrazoxane. In: StatPearls. StatPearls Publishing; 2024.

4. Deng S, Yan T, Jendrny C, et al. Dexrazoxane may prevent doxorubicin-induced DNA damage via depleting both topoisomerase II isoforms. BMC Cancer. 2014;14:842. doi:10.1186/1471-2407-14-842

5. Ichikawa Y, Ghanefar M, Bayeva M, et al. Cardiotoxicity of doxorubicin is mediated through mitochondrial iron accumulation. J Clin Invest. 2014;124(2):617–630. doi:10.1172/JCI72931

6. Osataphan N, Phrommintikul A, Chattipakorn SC, Chattipakorn N. Effects of doxorubicin-induced cardiotoxicity on cardiac mitochondrial dynamics and mitochondrial function: Insights for future interventions. J Cell Mol Med. 2020;24(12):6534–6557. doi:10.1111/jcmm.15305

7. Imaralu O, Singla D. Doxorubicin induces necroptosis in young mice. FASEB J. 2022;36(S1). doi:10.1096/fasebj.2022.36.S1.R4542

8. Abe K, Ikeda M, Ide T, et al. Doxorubicin causes ferroptosis and cardiotoxicity by intercalating into mitochondrial DNA and disrupting Alas1-dependent heme synthesis. Sci Signal. 2022;15(758):eabn8017. doi:10.1126/scisignal.abn8017

9. Linders AN, Dias IB, López Fernández T, Tocchetti CG, Bomer N, Van der Meer P. A review of the pathophysiological mechanisms of doxorubicin-induced cardiotoxicity and aging. npj Aging. 2024;10(1):9. doi:10.1038/s41514-024-00135-7

10. Murabito A, Hirsch E, Ghigo A. Mechanisms of Anthracycline-Induced Cardiotoxicity: Is Mitochondrial Dysfunction the Answer? Front Cardiovasc Med. 2020;7:35. doi:10.3389/fcvm.2020.00035

11. Xia P, Liu Y, Cheng Z. Signaling pathways in cardiac myocyte apoptosis. Biomed Res Int. 2016;2016:9583268. doi:10.1155/2016/9583268

12. Fiedler LR, Chapman K, Xie M, et al. MAP4K4 Inhibition Promotes Survival of Human Stem Cell-Derived Cardiomyocytes and Reduces Infarct Size In Vivo. Cell Stem Cell. 2019;24(4):579–591.e12. doi:10.1016/j.stem.2019.01.013

13. Te Lintel Hekkert M, Newton G, Chapman K, et al. Preclinical trial of a MAP4K4 inhibitor to reduce infarct size in the pig: does cardioprotection in human stem cell-derived myocytes predict success in large mammals? Basic Res Cardiol. 2021;116(1):34. doi:10.1007/s00395-021-00875-7

14. Wu Y, Xiu W, Wu Y. Salvianolic Acid A Protects H9C2 Cardiomyocytes from Doxorubicin-Induced Damage by Inhibiting NFKB1 Expression Thereby Downregulating Long-Noncoding RNA (lncRNA) Plasmacytoma Variant Translocation 1 (PVT1). Med Sci Monit. 2021;27:e929824. doi:10.12659/MSM.929824

15. Wen Y, Shen F, Wu H. Role of C5a and C5aR in doxorubicin-induced cardiomyocyte senescence. Exp Ther Med. 2021;22(4):1114. doi:10.3892/etm.2021.10548

16. Moulin M, Piquereau J, Mateo P, et al. Sexual dimorphism of doxorubicin-mediated cardiotoxicity: potential role of energy metabolism remodeling. Circ Heart Fail. 2015;8(1):98–108. doi:10.1161/CIRCHEARTFAILURE.114.001180

17. Deci MB, Ferguson SW, Scatigno SL, Nguyen J. Modulating Macrophage Polarization through CCR2 Inhibition and Multivalent Engagement. Mol Pharm. 2018;15(7):2721–2731. doi:10.1021/acs.molpharmaceut.8b00237

18. Wang J, Lee CJ, Deci MB, et al. MiR-101a loaded extracellular nanovesicles as bioactive carriers for cardiac repair. Nanomedicine. 2020;27:102201. doi:10.1016/j.nano.2020.102201

19. Wang J, Seo MJ, Deci MB, Weil BR, Canty JM, Nguyen J. Effect of CCR2 inhibitor-loaded lipid micelles on inflammatory cell migration and cardiac function after myocardial infarction. Int J Nanomedicine. 2018;13:6441–6451. doi:10.2147/IJN.S178650

20. Jasiewicz NE, Mei K-C, Oh HM, et al. Zippercells exhibit enhanced accumulation and retention at the site of myocardial infarction. Adv Healthc Mater. 2023;12(4):e2201094. doi:10.1002/adhm.202201094

21. Jasiewicz NE, Mei K-C, Oh HM, et al. In situ-crosslinked Zippersomes enhance cardiac repair by increasing accumulation and retention. Bioeng Transl Med. 2024;9(6):e10697. doi:10.1002/btm2.10697

22. Adams RA, Liu Z, Hsieh C, et al. Structural Analysis of Mitochondria in Cardiomyocytes: Insights into Bioenergetics and Membrane Remodeling. Curr Issues Mol Biol. 2023;45(7):6097–6115. doi:10.3390/cimb45070385

23. Larhammar M, Huntwork-Rodriguez S, Rudhard Y, Sengupta-Ghosh A, Lewcock JW. The Ste20 Family Kinases MAP4K4, MINK1, and TNIK Converge to Regulate Stress-Induced JNK Signaling in Neurons. J Neurosci. 2017;37(46):11074–11084. doi:10.1523/JNEUROSCI.0905-17.2017

24. Fu HY, Sanada S, Matsuzaki T, et al. Chemical Endoplasmic Reticulum Chaperone Alleviates Doxorubicin-Induced Cardiac Dysfunction. Circ Res. 2016;118(5):798–809. doi:10.1161/CIRCRESAHA.115.307604

25. Johnston RA, Rawling T, Chan T, Zhou F, Murray M. Selective inhibition of human solute carrier transporters by multikinase inhibitors. Drug Metab Dispos. 2014;42(11):1851–1857. doi:10.1124/dmd.114.059097

26. Loos M, Klampe B, Schulze T, et al. Human model of primary carnitine deficiency cardiomyopathy reveals ferroptosis as a novel mechanism. Stem Cell Reports. 2023;18(11):2123–2137. doi:10.1016/j.stemcr.2023.09.002

27. Windak R, Müller J, Felley A, et al. The AP-1 transcription factor c-Jun prevents stress-imposed maladaptive remodeling of the heart. PLoS ONE. 2013;8(9):e73294. doi:10.1371/journal.pone.0073294

28. Su L, Zhang G, Jiang L, Chi C, Bai B, Kang K. The role of c-Jun for beating cardiomycyte formation in prepared embryonic body. Stem Cell Res Ther. 2023;14(1):371. doi:10.1186/s13287-023-03544-9

29. Schumacher D, Curaj A, Staudt M, et al. Phosphatidylserine supplementation as a novel strategy for reducing myocardial infarct size and preventing adverse left ventricular remodeling. Int J Mol Sci. 2021;22(9). doi:10.3390/ijms22094401

30. Harel -Adar T, Ben Mordechai T, Amsalem Y, Feinberg MS, Leor J, Cohen S. Modulation of cardiac macrophages by phosphatidylserine-presenting liposomes improves infarct repair. Proc Natl Acad Sci USA. 2011;108(5):1827–1832. doi:10.1073/pnas.1015623108

31. Zhang L, Li Y, Liu X, et al. Optimal development of apoptotic cells-mimicking liposomes targeting macrophages. J Nanobiotechnology. 2024;22(1):501. doi:10.1186/s12951-024-02755-3

