## Supplemental Information for "Phosphatidylserine-Based Liposomes Encapsulating DMX-5804 Protect Against Doxorubicin-Induced Cardiotoxicity"

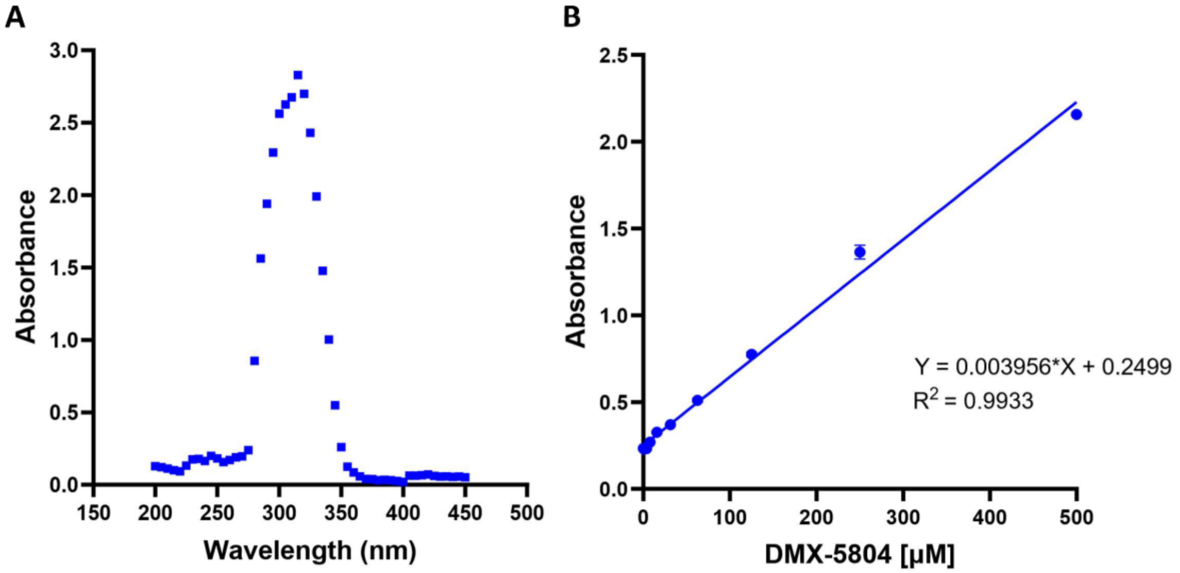


**Supplemental Figure 1:** Determination of λ max and calibration curve for DMX-5804 quantification. (A) UV–visible absorbance spectrum of DMX-5804 measured across the indicated wavelength range to identify the wavelength of maximum absorbance (λ max), which was used for all subsequent quantitative measurements. (B) Standard calibration curve generated by measuring absorbance at λ max across increasing concentrations of DMX-5804. A linear relationship between absorbance and concentration was observed over the range tested, and the least-squares linear regression (equation and $R^2$ shown on plot) was used for quantitative determination of DMX-5804 concentrations in downstream experiments. Data are shown as mean ± SD where applicable.

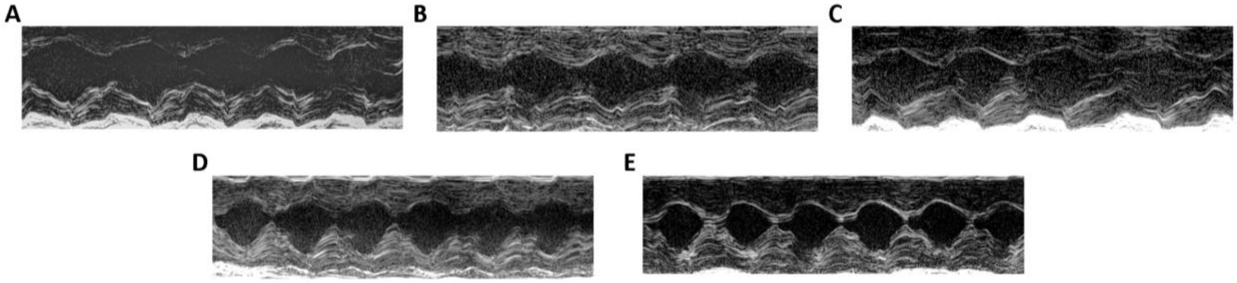


**Supplemental Figure 2:** Representative M-mode echocardiograms acquired on day 10 following treatment are shown for animals receiving (A) PBS vehicle control, (B) PS-Free NanoDMX Vehicle, (C) PS-Free NanoDMX, (D) NanoDMX Vehicle, and (E) NanoDMX. Images are presented to illustrate qualitative differences in left ventricular wall motion and chamber dynamics across dosing groups. Quantitative echocardiographic measurements derived from these recordings are reported separately.

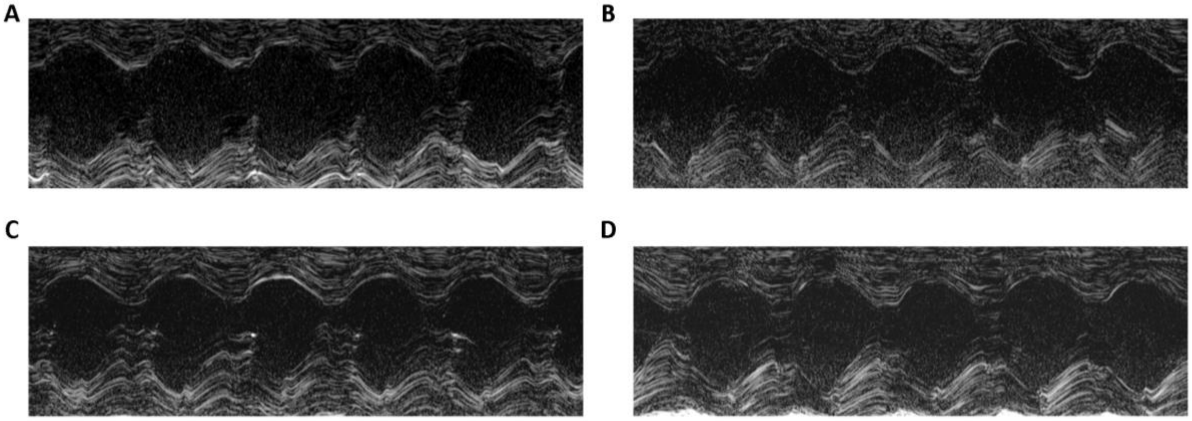


**Supplemental Figure 3:** Representative M-mode echocardiograms acquired on day 10 following treatment are shown for animals receiving (A) PBS vehicle control, (B) 25 mg/kg, (C) 35 mg/kg, and (D) 45 mg/kg of NanoDMX. Images are presented to illustrate qualitative differences in left ventricular wall motion and chamber dynamics across dosing groups. Quantitative echocardiographic measurements derived from these recordings are reported separately.

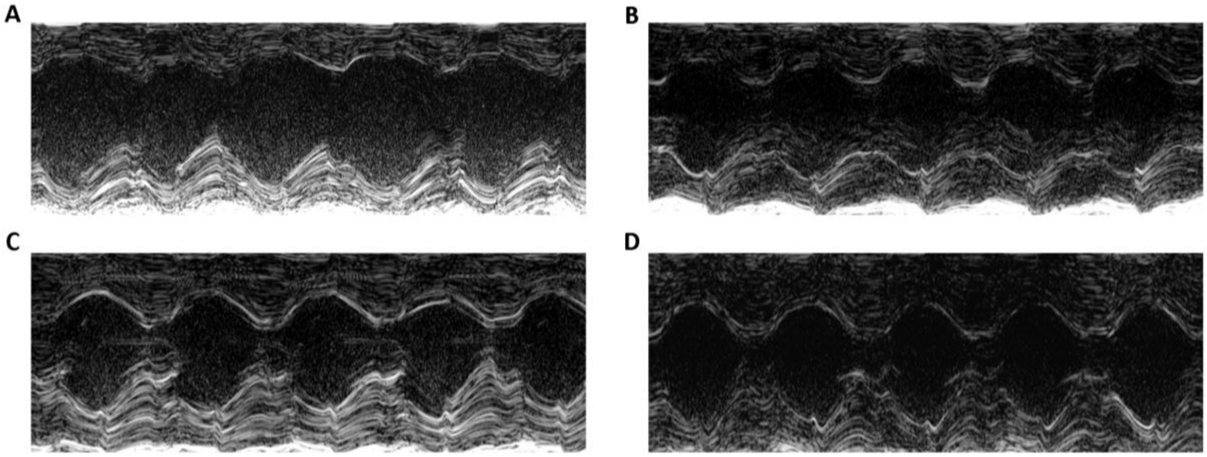


**Supplemental Figure 4:** Representative M-mode echocardiograms acquired on day 10 following treatment are shown for animals receiving (A) PBS vehicle control, (B) orally administered DMX-5804, (C) intravenously administered NanoDMX vehicle, and (D) intravenously administered NanoDMX. Images are presented to illustrate qualitative differences in left ventricular wall motion and chamber dynamics across dosing groups. Quantitative echocardiographic measurements derived from these recordings are reported separately.

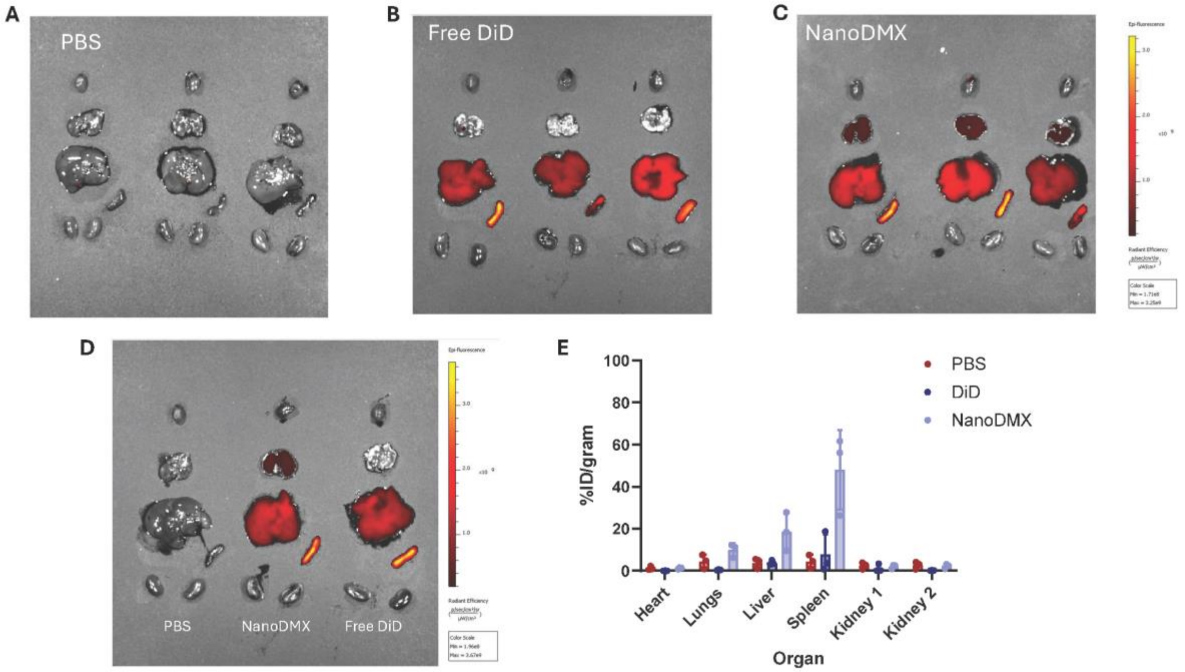


**Supplemental Figure 5:** NanoDMX liposomes were labeled with DiD (0.1 mol%) and administered intravenously following intraperitoneal doxorubicin (20 mg/kg). Free DiD and PBS served as controls. Blood was collected 5 min post-injection to determine the total injected dose. At 4 h post-injection, major organs were harvested and fluorescence was quantified by IVIS. Biodistribution is reported as percent injected dose per gram of tissue (%ID/g). (A) Ex vivo fluorescence images of organs from PBS-treated mice. (B) Ex vivo fluorescence images of organs from free DiD–injected mice. (C) Ex vivo fluorescence images of organs from NanoDMX-injected mice. (D) Representative ex vivo organ images from one mouse per treatment group. (E) Quantitative organ biodistribution expressed as %ID/g. Data represent mean and standard deviation from n = 3 mice per group. Fluorescence was quantified using identical IVIS acquisition settings across all samples. Left and right kidneys were quantified separately. Blood fluorescence was used to calculate the total injected dose for normalization.

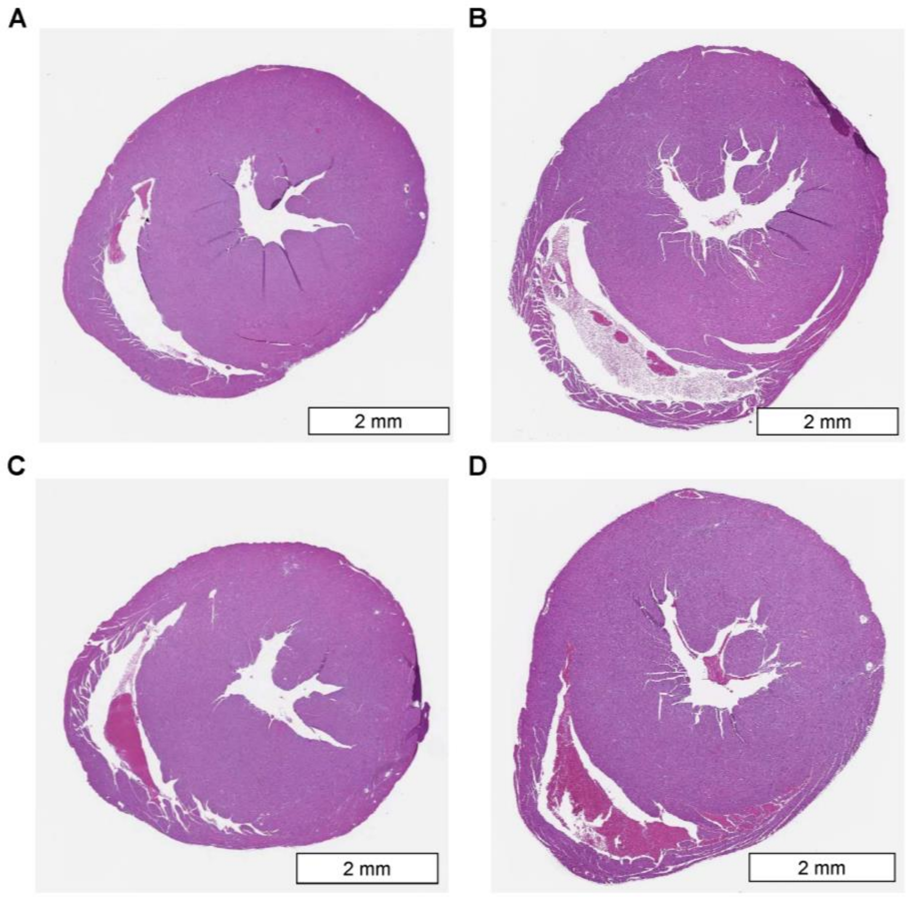


**Supplemental Figure 6:** Representative low-magnification histological sections of mouse hearts following doxorubicin treatment and NanoDMX treatment. Representative transverse cardiac sections imaged at 1.6× magnification are shown for (A) healthy control mice, (B) doxorubicin-treated mice, (C) doxorubicin-treated mice receiving oral DMX-5804, and (D) doxorubicin-treated mice receiving NanoDMX. Sections illustrate gross cardiac morphology and myocardial architecture across treatment groups. Scale bars represent 2 mm.
